# ATP hydrolysis by the SNF2 domain of the ultraspecific maintenance methylase Dnmt5 drives recognition and modification of hemimethylated DNA

**DOI:** 10.1101/2020.01.05.895227

**Authors:** Phillip A. Dumesic, Caitlin I. Stoddard, Sandra Catania, Geeta J. Narlikar, Hiten D. Madhani

**Affiliations:** Department of Biochemistry and Biophysics, University of California, San Francisco, CA 94158, USA

**Author notes:** To whom correspondence should be addressed. (H.D.M.); (G.J.N.).

**Keywords:** DNA methylation, DNA methyltransferase, ATPase, Epigenetics, Dnmt5, Enzyme mechanism

## Abstract

*C. neoformans* Dnmt5 is an ultraspecific maintenance-type CpG methyltransferase (DNMT) that mediates long-term epigenome evolution. It harbors a DNMT domain and SNF2 ATPase domain. We find that the SNF2 domain couples substrate specificity to an ATPase step that is essential for DNA methylation. Such coupling occurs independently of nucleosomes. Hemimethylated DNA preferentially stimulates ATPase activity, and mutating the Dnmt5 ATP binding pocket disproportionately reduces ATPase stimulation by hemimethylated versus unmethylated substrates. Engineered DNA substrates that stabilize a reaction intermediate by mimicking a ‘flipped-out’ conformation of the target cytosine bypass the SNF2 domain’s requirement for hemimethylation. This result implies that ATP hydrolysis by the SNF2 domain is coupled to base-flipping by the DNMT domain. These findings establish a new role for a SNF2 ATPase domain: controlling substrate recognition and catalysis by an adjoined enzymatic domain. This coupling may contribute to the exquisite specificity of Dnmt5 via mechanisms related to kinetic proofreading.

## INTRODUCTION

Methylation of the C5 position of cytosine (5mC) in eukaryotic genomes is a means to mark DNA sites in a potentially heritable manner. 5mC is present at repetitive elements and transposons, where it silences these elements for genome defense (Law and Jacobsen, 2010; Suzuki and Bird, 2008). Methylation is also found in the bodies of active genes, where its purpose is less clear (Feng et al., 2010; Huff and Zilberman, 2014; Schübeler, 2015; Zemach et al., 2010). In vertebrates, cytosine methylation is more widespread and occurs primarily at CG sites, both in gene bodies and intergenic regulatory regions. It is required for mammalian development, and can enforce long-term transcriptional repression of targeted genes, for instance during X chromosome inactivation, genomic imprinting, transposon suppression, and lineage-specific gene silencing (Jaenisch and Bird, 2003; Li et al., 1993; Smith and Meissner, 2013; Velasco et al., 2010).

An important aspect of cytosine methylation is its potential to enable the inheritance of gene silencing after loss of the initiating signal (Jones and Liang, 2009). In the simplest model, 5mC is maintained over cell divisions by a ‘maintenance’ DNA methyltransferase that acts preferentially on hemimethylated CG sites produced during DNA replication (Holliday and Pugh, 1975; Riggs, 1975). In contrast, an ‘establishment’ DNA methyltransferase deposits the initial 5mC mark de novo on an unmethylated CG template. Mammals encode DNA methyltransferases that broadly fall into these two classes. DNMT1 is widely expressed, is recruited to chromatin during DNA replication, and shows a kinetic preference for activity on hemimethylated as opposed to unmethylated substrates (Jeltsch and Jurkowska, 2016). In contrast, DNMT3A and DNMT3B exhibit equivalent activities on unmethylated and hemimethylated substrates, and are highly expressed in embryonic periods during which DNA methylation patterns are established.

Despite these general trends, high-resolution DNA methylation data have suggested that the segregation of roles between DNMT3A/B and DNMT1 might not be absolute: both enzyme classes appear to play a measurable role in DNA methylation establishment as well as maintenance (Arand et al., 2012; Jeltsch and Jurkowska, 2014; Jones and Liang, 2009; Riggs and Xiong, 2004). These results are consistent with the fact that the DNMT1 in vitro catalytic preference for hemimethylated sites is only 30-40-fold, seemingly insufficient for DNMT1 to faithfully preserve 5mC marks in a maintenance-only fashion, since this would require faithful replication of each of the 56 million CG sites in the human genome as either unmethylated or hemimethylated (Jeltsch, 2006; Jeltsch and Jurkowska, 2014). Therefore, ongoing de novo DNA methylation has been suggested to explain the stability of many 5mC marks. These findings highlight how the enzymological properties of an organism’s DNA methyltransferases might impose restrictions on how long the 5mC mark can be epigenetically inherited after loss of the mark’s initiating signal.

In this regard, our recent studies of the Dnmt5 family of eukaryotic cytosine methyltransferases are relevant. In this family—which is represented in fungi as well as in chlorophyte algae and stramenopiles—the DNMT domain is embedded in a multidomain architecture that includes a C-terminal SNF2 helicase-like domain (Iyer et al., 2011; Ponger and Li, 2005). Loss of function genomic studies revealed that Dnmt5 family methyltransferases are responsible for cytosine methylation at CG sites, and in some organisms represent the sole enzyme responsible for CG methylation (Huff and Zilberman, 2014). Our recent characterization of Dnmt5 in one such organism, the yeast *Cryptococcus neoformans*, revealed that the purified enzyme is an exquisitely specific maintenance type methyltransferase in vitro with no detectable de novo activity. Consistent with this observation, Dnmt5 was unable to restore 5mC landscapes when deleted from and subsequently reintroduced into a *C. neoformans* strain. Phylogenetic and functional analyses revealed that a species ancestral to *Cryptococcus* encoded a second DNA methyltransferase, DnmtX, that possesses de novo methyltransferase activity, but this gene was lost 50-150 million years ago. Further work demonstrated that cytosine methylation has been maintained for millions of years since this loss through a process analogous to Darwinian evolution in which methylation persistence requires epigenetic inheritance, rare random 5mC losses and gains, and natural selection (Catania et al., in press).

Here, we biochemically characterize *C. neoformans* Dnmt5 and uncover functional couplings between its SNF2 and DNA methyltransferase domains that may contribute to its unusually high specificity for hemimethylated substrates. We find that ATP hydrolysis by the SNF2 domain is essential for DNA methyltransferase activity by the DNMT domain. Non-hydrolyzable nucleotide analogs perturb Dnmt5’s affinity for DNA substrates and block DNA methylation activity. Strikingly, engineered DNA substrates predicted to stabilize the DNMT domain at intermediate steps of the cytosine methylation precatalytic pathway are sufficient to fully stimulate ATPase activity. These results demonstrate that SNF2-mediated ATP hydrolysis is both a response to recognition of a hemimethylated CG substrate and also itself required for productive substrate methylation. We discuss the possibility that these properties allow kinetic proofreading to enhance Dnmt5 specificity for its preferred substrates. More generally, these results reveal a previously undescribed role for a SNF2 ATPase in a DNA modification enzyme.

## RESULTS

### Dnmt5 is an ATP-dependent cytosine methyltransferase with high specificity for hemimethylated substrates

Dnmt5 proteins have unique architecture among DNA methyltransferase families: they contain a highly diverged DNMT domain followed by a RING domain, and a C-terminal region with SNF2 family homology (Huff and Zilberman, 2014; Iyer et al., 2011; Ponger and Li, 2005) (Figure 1A). Dnmt5 orthologs mediate CG methylation in diverse eukaryotes, including the yeast *C. neoformans*, where Dnmt5 is the organism’s sole DNA methyltransferase. Initial characterization of Dnmt5 in this system revealed it to have a strong in vivo and in vitro preference for action on hemimethylated, but not unmethylated, DNA substrates, and that it required ATP addition to the reaction (Catania et al., in press). The unusual specificity profile of this enzyme motivated us to examine the regulation of its activity in vitro, and to interrogate the role of its SNF2 domain.

**Figure 1.**
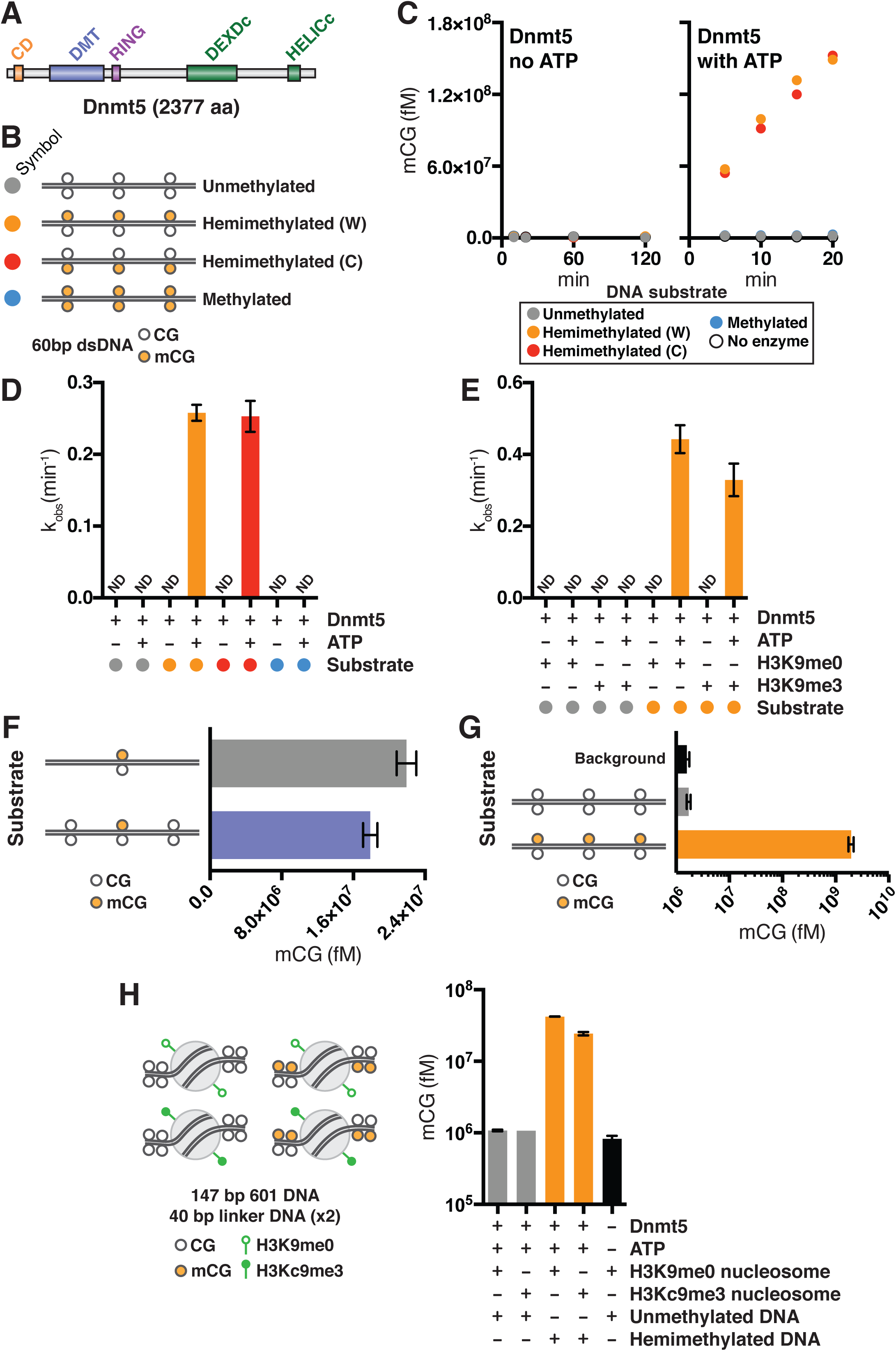
Dnmt5 is an ATP-dependent DNA methyltransferase with high specificity for hemimethylated substrates. **(A)** Predicted protein domains in *C. neoformans* Dnmt5. **(B)** dsDNA substrates used in methyltransferase experiments. Each 60bp substrate contains three CG sites that are either unmethylated, hemimethylated, or symmetrically methylated. **(C)** Example DNA methylation kinetics using 30 nM Dnmt5 and 5 μM each of the DNA substrates in B, with or without 1 mM ATP. **(D)** Average initial rates of Dnmt5 activity on DNA substrates described in B, in the presence or absence of ATP. ND: no detectable activity; error bars represent SD; n = 4-5. **(E)** Average initial rates of Dnmt5 activity on DNA substrates described in B, in the presence or absence of ATP and 5 μM histone tail peptides H3K9me0 or H3K9me3. ND: no detectable activity; error bars represent SD; n = 2-4. **(F)** Endpoint DNA methyltransferase activity of Dnmt5 (100 nM) on DNA substrates (50 nM) containing a single hemimethylated CG site as well as either zero or two unmethylated CG sites. Graph represents average and SD; n = 3. **(G)** Endpoint DNA methyltransferase activity of Dnmt5 (100 nM) on DNA substrates (5 μM) that are either unmethylated or hemimethylated. Background signal was measured in a reaction with unmethylated DNA but no enzyme. Graph represents average and SD; n = 3. **(H)** Endpoint DNA methylation activity of Dnmt5 (150 nM) on nucleosomal substrates (50 nM). For each substrate, DNA is 227 bp sequence corresponding to the Widom 601 nucleosome positioning sequence flanked on either side by a 40 bp linker sequence. Linkers each contain two CG sites that are either hemimethylated or unmethylated; the Widom 601 sequence is entirely unmethylated. Nucleosomes are either wild-type (H3K9me0) or MLA (H3Kc9me3). Graph represents averages and SD; n = 4.

We expressed the *C. neoformans* Dnmt5 ortholog in *S. cerevisiae* and isolated it to >90% purity (Figure S1A). Its DNA methyltransferase activity was tested under multiple turnover conditions using dsDNA substrates whose three CG motifs were either unmethylated, hemimethylated, or symmetrically methylated (Figure 1B). Confirming our prior results, in the absence of ATP, no activity was observed on any substrate. When ATP was added, activity was observed on hemimethylated substrates but not on unmethylated or symmetrically methylated substrates (Figure 1C). Dnmt5 exhibited an equivalent methyltransferase rate when acting on either the Watson or Crick strand of its hemimethylated substrate, and this rate was comparable to that of the well-studied maintenance methyltransferase DNMT1 (Pradhan et al., 1999) (Figure 1D). Importantly, when Dnmt5 was expressed and purified from its native host *C. neoformans* instead of from the orthologous system *S. cerevisiae*, protein yields were substantially lower, but the enzyme exhibited comparable qualitative and quantitative properties, confirming that its observed substrate preferences and ATP dependence were not an artifact of its expression system (Figure S1).

Given its lack of activity on unmethylated DNA substrates, we tested Dnmt5 under conditions that might unveil a latent de novo methyltransferase activity. First, since the Dnmt5 chromodomain has been shown to recognize trimethylation of histone H3 at lysine 9 (Catania et al., in press), we tested Dnmt5 activity in the presence of peptides corresponding to the histone H3 N-terminal tail, with or without K9 trimethylation. When saturating concentrations of either peptide were present, reaction rates on hemimethylated substrates were comparable, and activity remained undetectable on unmethylated DNA (Figure 1E). Second, since DNMT1 activity on unmethylated CG motifs can be stimulated by nearby methylated CG motifs (Tollefsbol and Hutchison, 1997), we tested whether the presence of a single hemimethylated CG site would unveil Dnmt5 activity on nearby unmethylated CG sites. To do so, an excess concentration of Dnmt5 was incubated with a DNA substrate and methylation was assessed at reaction endpoint. Substrates that contained unmethylated CG sites in addition to a single hemimethylated CG site were not methylated more so than substrates without the unmethylated CG sites, suggesting no Dnmt5 de novo methyltransferase activity (Figure 1F). Third, we tested whether Dnmt5 activity on unmethylated DNA substrates could be detected under multiple turnover conditions after reactions of longer duration (4 hr) and increased concentration of the methyl donor ^3^H-SAM (8 μM). Tritium incorporation into the hemimethylated substrate was ∼1,200-fold greater than background level, as assessed in a reaction containing substrate DNA and ^3^H-SAM but no enzyme (Figure 1G). Tritium incorporation into the unmethylated substrate, however, was indistinguishable from background. Finally, Dnmt5 was examined under single turnover conditions in the presence of mononucleosomal substrates designed to more faithfully mimic the chromatin context of in vivo Dnmt5 activity. We reconstituted mononucleosomes that were flanked on either side by 40 bp linker DNA containing two CG sites, which were either unmethylated or hemimethylated (Figure 1H). We also generated analogous mononucleosomes that were modified using methyl-lysine analog technology to contain a histone mark associated with heterochromatic sites of Dnmt5 activity in vivo (denoted H3Kc9me3). Analogous to the case of non-nucleosomal DNA substrates, Dnmt5 was active on mononucleosomal substrates containing hemimethylated DNA, but not on substrates containing unmethylated DNA (Figure 1H). Furthermore, the H3Kc9me3 mark did not unveil activity on unmethylated substrates (Figure 1H).

### Dnmt5 ATPase activity is sensitive to DNA substrate methylation and is required for cytosine methylation

We next investigated the nature of ATP’s requirement in DNA methylation by Dnmt5. When Dnmt5 was incubated with hemimethylated DNA, both Mg^2+^ and ATP were required for DNA methylation activity (Figure 2A). Dnmt5 was not active in the presence of ADP or any nucleotide analog tested, including non-hydrolyzable ATP analogs such as AMP-PNP (Figure 2B). Furthermore, AMP-PNP added to reactions containing Dnmt5 and ATP was able to compete with ATP and block DNA methylation (Figure S2A). These results suggested that ATP hydrolysis might be required for methyltransferase activity, a potential role for Dnmt5’s SNF2 family helicase-related domain. This domain belongs to a subfamily of RING finger-containing SNF2 domains, which are present in Rad5, Rad16, HLTF, and SHPRH (Flaus et al., 2006; Huff and Zilberman, 2014).

**Figure 2.**
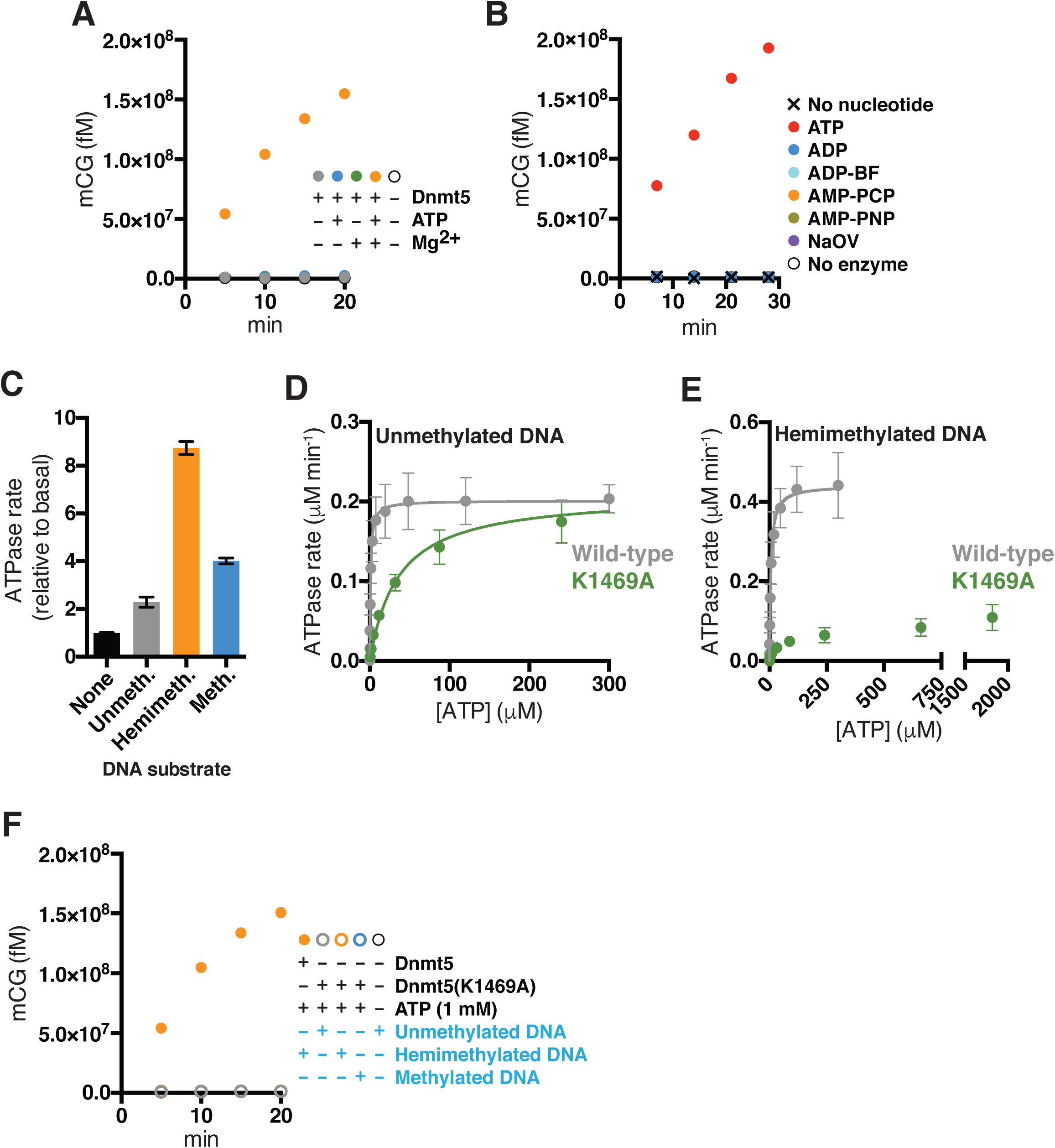
SNF2-mediated ATPase activity by Dnmt5 is sensitive to DNA substrate methylation and is required for DNA methyltransferase activity. (**A**) Example DNA methylation kinetics using 30 nM Dnmt5 and 5 μM hemimethylated DNA substrate, in the presence or absence of Mg^2+^ and ATP. (**B**) Example DNA methylation kinetics using 30 nM Dnmt5 and 5 μM hemimethylated DNA substrate, in the presence of ATP, ADP, or various nucleotide analogs: ADP beryllium fluoride (ADP-BP), AMP-PCP, AMP-PNP, or sodium orthovanadate (NaOV). (**C**) Average rates of ATPase activity in the presence of 40 nM Dnmt5 and 5 μM of the DNA substrates described in Figure 1B. Data are normalized to ATPase rate in absence of DNA (1 min^-1^). Error bars represent SD; n = 4. (**D-E**) Initial ATPase rates of 40 nM Dnmt5 or Dnmt5(K1469A) were determined at varying ATP concentrations in the presence of fixed, saturating concentrations of unmethylated (**D**) or hemimethylated (**E**) DNA substrates. Values were fitted to the Michaelis-Menten equation. Error bars represent SD; n = 4. (**F**) Example DNA methylation kinetics using 30 nM Dnmt5 or Dnmt5(K1469A) and 5 μM of the DNA substrates described in Figure 1B.

To test whether Dnmt5 hydrolyzes ATP, we performed an NADH-coupled ATPase assay using full-length Dnmt5. Dnmt5 exhibited a basal ATPase activity of approximately 1 min^-1^, which was stimulated 2-fold by saturating concentrations of unmethylated dsDNA (Figure 2C). Unmethylated dsDNA substrates of various lengths stimulated ATPase activity, and the presence of CG sites was not required (Figure S2B,C). We next tested whether hemimethylated DNA would differentially stimulate ATPase activity. Indeed, when Dnmt5 was incubated with saturating concentrations of 60 bp dsDNA substrates that contained three CG sites differing in their methylation status (unmethylated, hemimethylated, or symmetrically methylated), distinct ATPase stimulation effects were observed (Figure 2C). Hemimethylated DNA was most stimulatory, whereas symmetrically methylated DNA was intermediate and unmethylated DNA least stimulatory. Measurement of *K_m,app_*^DNA^ for unmethylated and hemimethylated substrates demonstrated that both *K_m_* and *k*_cat_ effects contribute to the preferential ATPase stimulation by hemimethylated DNA (Figure S2D). The efficacy with which hemimethylated substrates stimulated ATPase activity did not depend on their length or on the number of hemimethylated CG sites they contained (Figure S2E).

In a complementary set of experiments, we measured the *K_m,app_*^ATP^ and *k*_cat_ ^ATPase^ parameters by varying ATP concentration in the presence of saturating concentrations of unmethylated or hemimethylated DNA substrates. The parameters differed as a function of the DNA substrate, with hemimethylated DNA effecting significantly greater *K_m,app_*^ATP^ and *k*_cat_ ^ATPase^ than did unmethylated DNA (Table 1). These findings indicate that the methylation state of the DNA substrate bound to Dnmt5 alters interactions made within the nucleotide binding pocket of the SNF2 domain.

We next mutated the Walker A motif of the Dnmt5 SNF2 domain (K1469A) to confirm the SNF2 domain’s role in ATPase activity (Figure S2F). As expected given the Walker A motif’s role in ATP binding, this mutation increased *K_m,app_*^ATP^ but had no effect on k*_cat_* ^ATPase^ in the presence of unmethylated DNA substrates (Figure 2D and Table 1). Curiously, the mutation had a different effect on ATPase activity in the setting of hemimethylated DNA, where no tested concentration of ATP could fully rescue Dnmt5(K1469A) ATPase activity (Figure 2D and E). This result suggests an additional role—beyond simply ATP binding—for the K1469 residue when hemimethylated DNA is present. We reasoned that this unique role could involve coupling ATP hydrolysis to productive recognition of the hemimethylated substrate by the DNMT domain. We therefore tested the methyltransferase activity of Dnmt5(K1469A). Strikingly, Dnmt5(K1469A) showed no detectable methyltransferase activity even at a 1 mM ATP, a concentration at which its ATPase rate on hemimethylated substrates is readily detected (Figure 2E and F). Together, these results demonstrate that the SNF2 domain is responsive to DNA substrate methylation state, confirm that this domain is required for DNA methylation, and suggest that the K1469A mutation decouples ATPase activity from DNA methyltransferase activity.

### Dnmt5 binds DNA substrates with a preference for hemimethylation, and its affinity is modulated by nucleotide binding

We next assessed DNA substrate binding by Dnmt5 in order to determine whether this function is regulated by the ATP hydrolysis cycle of the SNF2 domain. We first purified an MBP-fused, truncated form of Dnmt5 containing only the DNMT domain (residues 345-747) and assessed its ability to bind DNA using an electrophoretic mobility shift assay (EMSA). When incubated with Cy5-labeled 60 bp DNA substrates corresponding to sequences used in the prior methyltransferase reactions, Dnmt5(345-747) bound unmethylated and hemimethylated DNA with similar affinity (K_d_ ∼ 7 nM) (Figure 3B). This result suggests that the DNMT domain by itself does not have binding specificity for the hemimethylated substrate. An MBP-fused Dnmt5 truncation containing only the N-terminal chromodomain (residues 1-150) did not show detectable DNA binding, confirming that the MBP tag was not responsible for DNA binding (Figure S3A). DNA binding could be competed by excess concentrations of unlabeled DNA (Figure S3B). As expected, Dnmt5(345-747), which lacks the SNF2 domain, did not exhibit DNA methyltransferase activity in vitro (Figure S3C).

**Figure 3.**
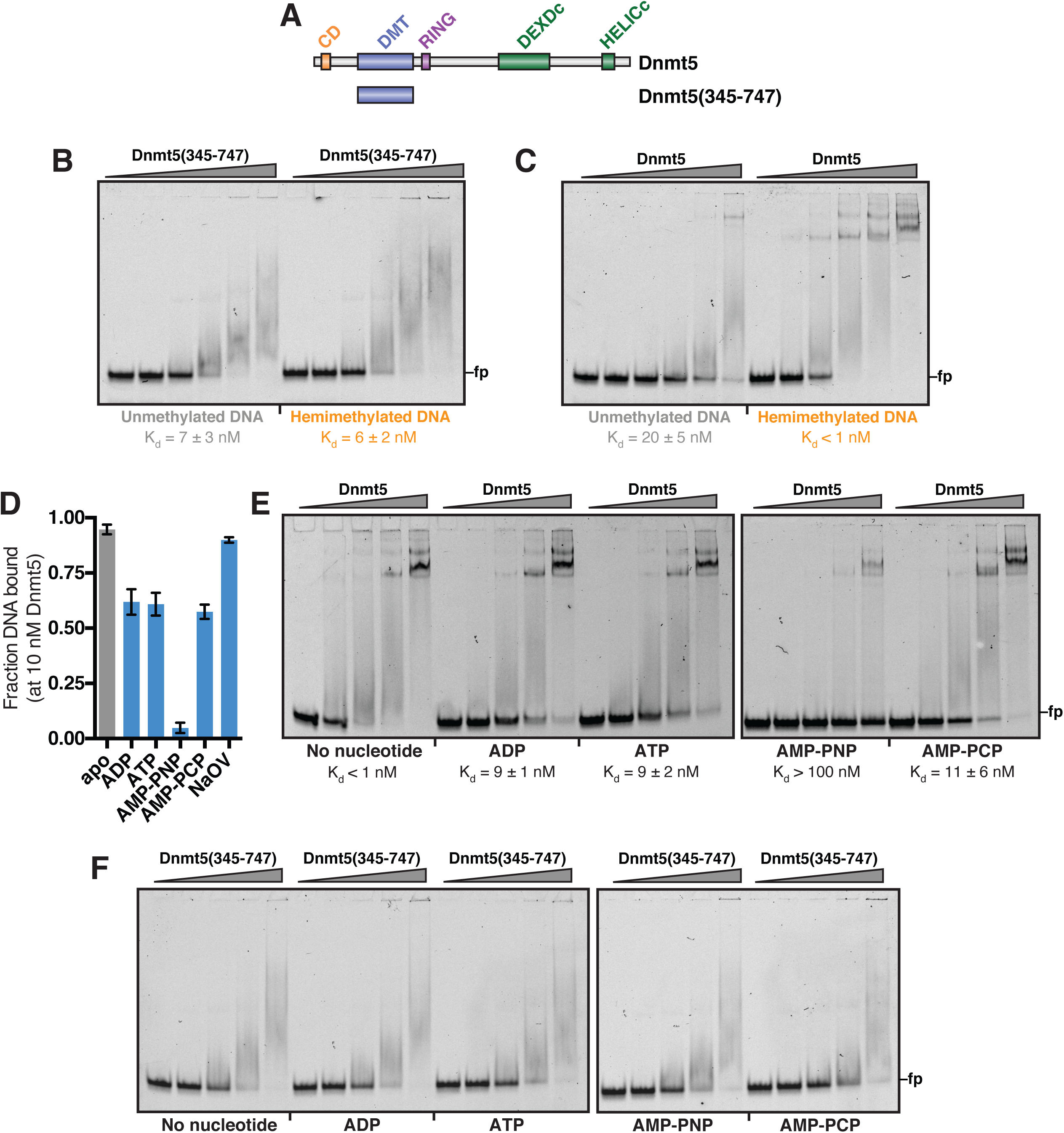
Nucleotide state of Dnmt5 modulates its DNA binding. (**A**) Schematic illustrating the Dnmt5(345-747) truncation. (**B-C**) EMSA assessing binding of Dnmt5(345-747) (0-67 nM) (**B**) or full-length Dnmt5 (**C**) to 1 nM labeled unmethylated or hemimethylated 60 bp dsDNA described in Figure 1B. K_d_ values represent average and SD of 3-5 experiments. (**D**) Screen of nucleotide analogs for effects on Dnmt5 DNA binding. Dnmt5 (10 nM) was incubated with labeled hemimethylated DNA in the presence of nucleotide or nucleotide analog (1 mM), and fraction probe bound was measured by EMSA. Graph represents average and SD; n = 4. (**E-F**) EMSA assessing binding of full-length Dnmt5 (0-50 nM) (**E**) or Dnmt5(345-747) (0-50 nM) (**F**) to 1 nM labeled hemimethylated DNA in the presence of nucleotide or nucleotide analog (1 mM). K_d_ values represent average and SD of 3 experiments. (**G**) Initial ATPase rates of 30 nM Dnmt5 were determined at varying concentrations of unmethylated or hemimethylated DNA substrates in the presence of fixed, saturating concentrations of ATP. Values were fitted to the Michaelis-Menten equation to determine *K_m,app_* ^DNA^, expressed as mean and standard error. Error bars represent SD; n = 3.

**Figure 4.**
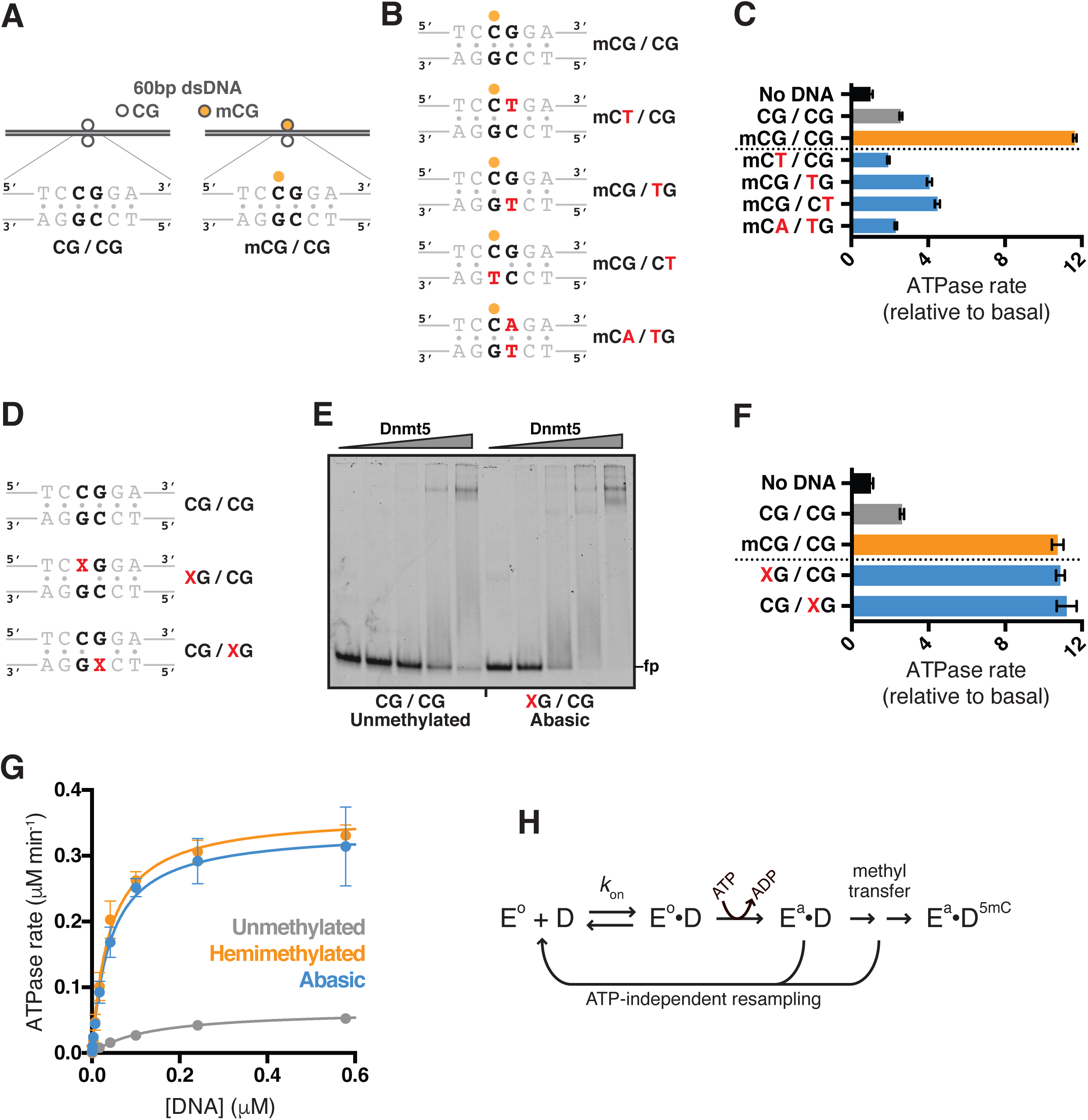
Dnmt5 ATPase activity is stimulated by DNA substrates predicted to stabilize intermediate steps of the cytosine methylation precatalytic pathway. (**A**) DNA substrates to assess effects of CG site manipulation. Each 60bp dsDNA substrate contains one CG site that is either unmethylated or hemimethylated. (**B**) Schematics of CG site mutant hemimethylated DNA substrates. Red color indicates base mutation as compared to original CG sequence. (**C**) Average rates of ATPase activity in the presence of 40 nM Dnmt5 and 5 μM of the DNA substrates described in panels A and B. Data are normalized to ATPase rate in absence of DNA (1 min^-1^). Error bars represent SD; n = 4. (**D**) Schematics of abasic DNA substrates. Red color indicates base mutation as compared to original CG sequence as in panel A. ‘X’ indicates abasic site. (**E**) EMSA assessing binding of full-length Dnmt5 (0-67 nM) to 1 nM labeled unmethylated or abasic site DNA. (**F**) Average rates of ATPase activity in the presence of 40 nM Dnmt5 and 5 μM of the DNA substrates described in panels A and E. Data are normalized to ATPase rate in absence of DNA (1 min^-1^). Error bars represent SD; n = 3-4. (**G**) Initial ATPase rates of 30 nM Dnmt5 or Dnmt5(K1469A) were determined at varying concentrations of DNA substrate (unmethylated, hemimethylated, or abasic site) in the presence of fixed, saturating concentrations of ATP. Values were fitted to the Michaelis-Menten equation. Error bars represent SD; n = 3. (**H**) Hypothetical model of ATPase stimulation by an intermediate step in the cytosine methylation precatalytic pathway. The inactive Dnmt5 enzyme (E°) binds DNA (**D**), leading to conformational changes that include cytosine flipping out of the DNA helix and activation loop closure. The resulting E•D complex is competent for ATPase activity, resulting in an increased population of E^a^, an active enzyme capable of proceeding in DNA methylation. Each indicated step—DNA binding, ATP hydrolysis, and methyltransfer—may be sensitive to the methylation state of the DNA substrate, conferring selectivity for maintenance DNA methylation.

In order to test whether the Dnmt5 N- and C-terminal regions (peripheral to DNMT domain) influence DNA binding (Figure 3A), we performed EMSAs using full-length Dnmt5. Unlike Dnmt5(345-747), full-length Dnmt5 bound DNA differentially depending on methylation state (Figure 3C). The K_d_ of Dnmt5 for unmethylated and hemimethylated DNA was 20 ± 5 nM and <1 nM, respectively, the latter being an upper bound due to limitations of our electrophoretic mobility shift assay (EMSA). Dnmt5 binding to labeled DNA could be competed with excess concentrations of unmethylated or hemimethylated unlabeled DNA, confirming assay specificity (Figure S3D). Thus, as compared to the isolated DNMT domain, the full-length Dnmt5 protein shows at least a 20-fold greater substrate binding specificity (i.e., its preference for hemimethylated versus unmethylated DNA).

The above results suggest that the N- and/or C-terminal domains present in full-length Dnmt5 help confer specificity for hemimethylated DNA by decreasing affinity for the unmethylated substrate and increasing affinity for the hemimethylated substrate. We hypothesized that the SNF2 domain plays such a role, and therefore tested whether manipulation of the SNF2 domain via nucleotides or nucleotide analogs would affect the ability of full-length Dnmt5 to bind hemimethylated DNA. Relative to the apo state, addition of ADP, ATP, AMP-PCP, and AMP-PNP reduced Dnmt5 affinity for hemimethylated DNA, with AMP-PNP having the largest effect (Figure 3D). We validated these effects by performing quantitative EMSA experiments in the presence of each nucleotide or analog (Figure 3E). Dissociation constants measured in this way were concordant with *K_m,app_*^DNA^ values for the same DNA substrates as measured by ATPase assay at equivalent salt concentrations (Figure S2D). Similar effects of nucleotides and analogs were observed in the context of Dnmt5 binding to unmethylated DNA (Figure S3E). Importantly, nucleotides had no effect on DNA binding by Dnmt5(345-747), since this truncation does not contain the SNF2 domain or its nucleotide-binding site (Figure 3F). These results indicate that the nucleotide-bound state of Dnmt5 influences its DNA binding ability, demonstrating a coupling between its SNF2 and DNMT domains.

### Dnmt5 ATPase activity is responsive to CG base manipulations

The selective responsiveness of Dnmt5 ATPase activity to hemimethylated DNA suggested that ATPase activity is coupled to detection of optimal substrates for the methyltransferase activity of Dnmt5. We therefore sought to more specifically define the DNA features that stimulate ATPase activity, with a focus on the hemimethylated CG site itself.

We generated 60 bp dsDNA substrates that contain a single CG site, which was either unmethylated (CG/CG) or hemimethylated (mCG/CG) (Figure 4A). Each base in the mCG/CG motif, excepting the methylated cytosine, was then individually mutated to thymine (Figure 4B). We also created a mutated substrate in which two alterations were made in order to preserve DNA base-pairing (mCA/TG). As expected, the hemimethylated substrate caused a greater stimulation of ATPase activity than did the unmethylated substrate. Each CG mutation reduced the ability of the DNA substrate to stimulate ATPase activity, even with the substrates present at saturating concentrations (Figure 4C). Therefore, unlike unmethylated DNA, which stimulates ATPase activity to the same modest extent in the presence or absence of CG motifs (Figure S2C), hemimethylated DNA strictly requires the intact CG motif for its full effect.

The sensitivity of ATPase activity to the CG motif raised the possibility that the SNF2 domain is coupled to hemimethylated CG detection by the DNMT domain, a protein fold able to make multiple contacts to a CG motif and to undergo conformational rearrangements when its preferred substrate is present (Matje et al., 2011; 2013; Song et al., 2012). To test this model, we sought a DNA substrate that would bind the DNMT domain and stabilize it in such a conformation. Extensive study of cytosine methylation by the M.HhaI methyltransferase has elucidated a precatalytic pathway in which the enzyme 1) binds its DNA substrate, 2) destabilizes the target cytosine, causing it to flip out of the DNA helix, and 3) undergoes a conformational rearrangement to close its catalytic loop, creating a ‘closed’ active site capable of methylation (Matje et al., 2011; Roberts and Cheng, 1998; Sankpal and Rao, 2002). Interestingly, replacing the target cytosine with an abasic site (while preserving the phosphate backbone and ribose ring) increases M.HhaI DNMT domain affinity for the substrate while stabilizing the ‘closed’ catalytic loop conformation (Matje et al., 2011; 2013; O’Gara et al., 1998). These findings have been attributed to the fact that absence of the target cytosine obviates the energetic cost of base flipping.

We predicted that such an abasic substrate would bind with increased affinity to Dnmt5 and stabilize its DNMT domain in a ‘closed’ catalytic loop conformation analogous to the intermediate step of the cytosine methylation precatalytic pathway. If progression through the DNMT domain precatalytic pathway is coupled to SNF2 domain activity, then this substrate would be expected to potently induce ATP hydrolysis, despite the fact that it does not contain a methylated cytosine. We generated 60 bp unmethylated dsDNA substrates in which either the Watson or Crick strand target cytosine was replaced by an abasic site (‘X’) (Figure 4D). As assessed by EMSA, the abasic substrate bound full-length Dnmt5 with 5-fold greater affinity than did the unmethylated substrate, consistent with the idea that this substrate engages the DNMT domain (Figure 4E). We next tested the ability of saturating concentrations of abasic DNA substrates to induce SNF2 domain ATPase activity. Remarkably, each abasic substrate, despite lacking a methylation mark, stimulated ATPase activity to the same extent as did the hemimethylated CG substrate (Figure 4F). Finally, to confirm that the abasic substrate fully recapitulated the ATPase stimulatory properties of the hemimethylated CG substrate, we measured *k*_cat_ and *K_m,app_*^DNA^ for each DNA substrate using an ATPase assay. The abasic substrate recapitulated the effects of the hemimethylated substrate (decreased *K_m,app_*^DNA^ and increased *k*_cat_), as compared to unmethylated DNA (Figure 4G). These results implicate the DNMT domain and its precatalytic conformational changes as inputs to SNF2-mediated ATPase activity.

## DISCUSSION

Our recent work indicates that Dnmt5 is an exceptionally specific maintenance methyltransferase both in vitro and in vivo. We therefore biochemically characterized its regulation and substrate specificity, and found this enzyme to exhibit a novel DNA methylation mechanism in which methyltransferase activity is coupled to the ATP hydrolysis activity of a SNF2 domain encoded on the same polypeptide. ATPase activity is responsive to hemimethylated CG motifs, and potentially regulated by DNMT domain conformational changes during the precatalytic pathway of DNA methylation. We discuss below the possibility that the coupling of DNMT activity to a seemingly needless expenditure of ATP enables increased Dnmt5 substrate specificity.

### Dnmt5 is a cytosine methyltransferase with high specificity for hemimethylated DNA substrates

The present findings extend our prior work demonstrating that Dnmt5 possesses extraordinary specificity for hemimethylated substrates. We detect no Dnmt5 in vitro methyltransferase activity on unmethylated CG sites in a variety of substrates, even in the context of nearby methylation marks or on nucleosomal substrates engineered to mimic the H3K9me3-decorated heterochromatin at the in vivo loci of Dnmt5 activity. Using endpoint DNA methylation assays under multiple-turnover conditions, we estimate ∼1000-fold as a lower bound of Dnmt5’s preference for hemimethylated DNA, which is probably a substantial underestimate of the true value given that Dnmt5 activity slows over the reaction course (likely owing to enzyme inhibition by accumulating levels of the reaction product SAH). Dnmt5’s specificity does not appear to require processive action, since it is observed using substrates containing only 1-3 hemimethylated CG sites. Furthermore, the fact that Dnmt5 reaction rates on hemimethylated substrates are similar to rates observed for DNMT1 argues that the observed specificity is not simply a result of grossly suboptimal reaction conditions.

Importantly, specificity was measured in the context of saturating levels of DNA substrates, and might be enhanced still further in vivo by the fact that Dnmt5 exhibits a binding affinity preference for hemimethylated DNA. The demonstrated recognition of H3K9 methylation by the Dnmt5 chromodomain could additionally contribute to specificity by favoring Dnmt5 localization at heterochromatic loci enriched in hemimethylated CG sites, as could the previously demonstrated roles of Uhrf1 and Swi6 in the Dnmt5 methylation system (Catania et al., in press). Although exact definitions of substrate specificity in DNA methyltransferases are challenging (Jeltsch, 2006), our observations indicate that Dnmt5 exhibits substantially more specificity than does the paradigmatic maintenance DNA methyltransferase DNMT1 (20-40-fold preference for hemimethylated DNA in vitro), and therefore appears to be unusually well-equipped to faithfully maintain 5mC marks, in a purely epigenetic fashion, over long timescales (Catania et al., in press; Jeltsch and Jurkowska, 2014).

### Dnmt5 ATP hydrolysis is stimulated by hemimethylated substrates and necessary for methyltransferase activity

The Dnmt5 family of DNA methyltransferases is characterized by the presence of a SNF2 helicase-like domain (Iyer et al., 2011; Ponger and Li, 2005), but the role of this domain—and its potential relationship to DNA methyltransferase activity—has not previously been investigated. Our present results demonstrate that SNF2-mediated ATP hydrolysis by Dnmt5 is strictly required for its DNA methyltransferase activity. Furthermore, this SNF2 activity is stimulated most effectively by the hemimethylated CG sites that represent optimal substrates for Dnmt5 DNA methyltransferase activity.

SNF2 homologs have long been implicated in DNA methylation. Lsh and DDM1 are required for DNA methylation in mouse and *Arabidopsis*, respectively (Dennis et al., 2001; Jeddeloh et al., 1999). The role of these factors appears distinct from that of Dnmt5, however. Lsh and DDM1 are thought to remodel nucleosomes in order to overcome the nucleosome’s inhibitory effect on DNA methylation (Brzeski and Jerzmanowski, 2003; Felle et al., 2011; Ren et al., 2015; Zemach et al., 2013). Indeed, the requirement for DDM1 in vivo can be suppressed by additional mutations (such as loss of histone H1) that impede nucleosome compaction (Zemach et al., 2013). In contrast, we find that the Dnmt5 SNF2 domain is absolutely required for DNA methylation even in non-nucleosomal contexts, arguing against nucleosome remodeling as its primary function. Furthermore, Dnmt5 belongs to a subfamily of SNF2 domains (with Rad5, HLTF, and SHPRH) that has been associated with resolution of DNA replication forks, but not with nucleosome remodeling activity (Huff and Zilberman, 2014; Unk et al., 2010).

For similar reasons, the role of the Dnmt5 SNF2 domain does not appear analogous to that of the SNF2 family member Mi-2, a component of the NuRD chromatin modifying complex (Allen et al., 2013; Torchy et al., 2015). Just as Dnmt5 comprises a DNA methyltransferase activity (by its DNMT domain) that requires ATP hydrolysis (by its SNF2 domain), NuRD comprises a histone deacetylase activity (by HDAC1/2) that requires SNF2-mediated ATP hydrolysis (by Mi-2) (Torchy et al., 2015). The requirement for ATP hydrolysis by Mi-2, however, can be bypassed when NuRD activity is assessed on histone octamers instead of nucleosomal DNA, indicating that the role of Mi-2 is to remodel nucleosomes in order to expose HDAC1/2 substrates (Verreault et al., 1998; Xue et al., 1998). This type of indirect mechanism cannot be posited for Dnmt5, since Dnmt5 requires ATP hydrolysis for DNA methylation even in the absence of nucleosomes. Furthermore, the presence of SNF2 and DNMT activities on the same Dnmt5 polypeptide (instead of on two distinct subunits of the NuRD chromatin-modifying complex) raises the possibility of direct enzymatic coupling between the two domains.

### ATP hydrolysis as a means for cytosine methyltransferase specificity

What, then, is the purpose of ATP hydrolysis by Dnmt5, an expenditure of energy not fundamentally required for the transfer of a methyl group from SAM to cytosine? A clue might come from our observations suggesting that not every ATP hydrolysis event leads to productive DNA methylation. Specifically, we found that the increased ability of hemimethylated substrates to stimulate ATPase activity (2-4-fold k_cat_^ATPase^ difference versus unmethylated substrates) was not nearly sufficient to explain the difference in DNA methyltransferase activity on these same two substrates (>1,000-fold). This discordance would not be possible if every ATPase hydrolysis event were linked to a productive DNA methylation event. It is therefore likely that steps of the DNA methylation reaction mechanism subsequent to ATP hydrolysis contribute additionally to substrate specificity. Another (non-mutually exclusive) possibility is that ATP hydrolysis in the context of unmethylated CG sites is ‘off pathway,’ and fundamentally not capable of promoting DNA methylation. This latter idea is underscored by the fact that no DNA methylation is detected on unmethylated substrates, despite appreciable Dnmt5 ATPase activity in this setting.

Several lines of evidence suggest that the presence of hemimethylated DNA stabilizes a conformation of the SNF2 ATPase domain distinct from its conformation in the presence of unmethylated DNA. First, the catalytic parameters of SNF2-mediated ATP hydrolysis are significantly different in these two conditions. Specifically, the *K_m,app_* ^ATP^ and k_cat_ ^ATPase^ are higher in the presence of hemimethylated substrate than in the presence of unmethylated substrate. Second, a Walker A site mutation (K1469A) has different effects on SNF2 activity in the presence of hemimethylated versus unmethylated substrates. In the presence of unmethylated DNA, this mutation’s effect can be rescued by high concentrations of ATP, consistent with a role for this residue in ATP binding. In contrast, in the presence of hemimethylated DNA, high ATP concentrations do not rescue SNF2 ATPase activity, and DNA methyltransferase activity is absent despite detectable ATPase activity. Therefore, the K1469 residue, uniquely in the setting of hemimethylated substrate, may play a role in addition to ATP binding, perhaps in coupling ATP hydrolysis to productive DNA methylation.

These observations are consistent with a model in which ATPase activity provides an opportunity for Dnmt5 to adopt an active conformation (E^a^ in Figure 4H) that is competent for DNA methylation. Conversely, ATPase activity is better stimulated by DNA substrates that stabilize the active conformation. Several aspects of this model could contribute to Dnmt5’s in vitro and in vivo preference for maintenance DNA methylation. First, Dnmt5 binds substrates with a preference for hemimethylated versus unmethylated DNA. Second, analogous to the behavior of ‘cognate’ substrates in the well-studied M.HhaI system, hemimethylated DNA substrates may more readily enable stabilization of the activated enzyme state, by virtue of target cytosine flipping and DNMT domain activation loop closure. In support of this idea, we find higher rates of ATP hydrolysis in the presence of hemimethylated substrates. Abasic substrates, which are expected to induce analogous DNMT domain conformational changes, likewise stimulate maximal ATPase activity. Third, after ATP hydrolysis, the chemical steps of DNA methylation may themselves be sensitive to DNA substrate methylation state.

Finally, we note that ATP hydrolysis in this model provides an extra, irreversible step between substrate binding and DNA methylation. Dissociation of the enzyme-substrate complex after hydrolysis, if occurring preferentially for unmethylated substrates as compared to hemimethylated substrates, would provide additional substrate specificity, since this discard step is thermodynamically driven out from the productive methylation pathway by virtue of ATP hydrolysis (Burgess and Guthrie, 1993; Hopfield, 1974; Ninio, 1975; Yarus, 1992a; 1992b). Our proposed model thus shares conceptual elements with classical models of kinetic proof-reading.

To our knowledge, Dnmt5 represents the first example of a reaction mechanism in which ATP hydrolysis is coupled to cytosine methylation. We find that the principal requirement of the Dnmt5 SNF2 domain is not nucleosome remodeling. Instead, we propose that ATP hydrolysis acts in concert with precatalytic events at the DNMT domain to provide an opportunity for increased substrate specificity. Consistent with such a coupling, our preliminary efforts to express truncated Dnmt5 constructs containing only the SNF2 domain resulted in proteins that did not bind DNA and could not be stimulated to hydrolyze ATP by any DNA substrate (data not shown). Admittedly, the idea that ATP hydrolysis contributes directly to DNMT domain substrate specificity is challenging to test owing to the complete lack of detectable DNA methyltransferase activity in the context of Dnmt5 mutants that do not hydrolyze ATP. The ability to design rationally more subtle mutations will require atomic resolution structures of Dnmt5 in its various states. Nevertheless, our model helps explain the remarkable ability of *C. neoformans* Dnmt5 to mediate epigenome evolution of million-year timescales in the absence of a de novo methyltransferase (Catania et al., in press).

## ACKNOWLEDGEMENTS

We thank C. Chio, D. Elnatan, L. Pack, J. Reuter, K. Verba, and members of the Madhani and Narlikar laboratories for helpful discussions, and N. Nguyen for media preparation.

P.A.D., C.I.S., G.J.N., and H.D.M. designed the study. P.A.D. and C.I.S. performed recombinant protein expression and purification. P.A.D. performed methyltransferase, ATPase, and EMSA assays. C.I.S. prepared nucleosomal substrates. S.C. generated *C. neoformans* strains. P.A.D., G.J.N., and H.D.M. wrote the manuscript. All authors contributed to editing the manuscript.

## EXPERIMENTAL PROCEDURES

### Recombinant protein expression and purification

Protein constructs were purified from *E. coli*, *C. neoformans*, or *S. cerevisiae*. For the former, a codon-optimized DNA sequence encoding the DNA methyltransferase domain of *C. neoformans* Dnmt5 (residues 345-747) was cloned into the pMAL vector. The *E. coli* strain BL21(DE3) was transformed, grown to OD_600_ = 0.8 in 2x YT medium, then induced with 1 mM IPTG overnight at 18°C. Recombinant MBP-Dnmt5(345-747)-6xHis was purified using Ni-NTA agarose resin (Qiagen) and measured by A_280_ (ε = 139,790 cm^-1^ M^-1^).

Using standard procedures (Chun and Madhani, 2010), a *C. neoformans* strain was generated in which the endogenous *DNMT5* gene was tagged (2xFLAG) and its promoter replaced by a galactose-inducible promoter (pGAL7) (Table S1). This strain was grown at 30°C to OD_600_ = 2.0 in 4 L YPAG medium (1% yeast extract, 2% Bacto-peptone, 2% galactose, 0.015% L-tryptophan, 0.004% adenine), at which point the cells were harvested, resuspended in TAP buffer (25 mM HEPES-KOH pH7.9, 0.1 mM EDTA, 0.5 mM EGTA, 2 mM MgCl_2_, 20% glycerol, 0.1% Tween-20, 300mM KCl, 1 mM DTT, 1x EDTA-free Complete protease inhibitor (CPI; Roche)), snap frozen, then lysed using a coffee grinder (3 min) and mortar and pestle (30 min). Lysate was resuspended in TAP buffer and centrifuged at 27,000 x *g* for 40 min at 4°C, after which it was incubated in batch format with anti-FLAG M2 affinity resin (Sigma) for 4 hr. The resin was washed three times with TAP buffer totaling 1 hr. Tagged protein was eluted by three washes at 4°C in FLAG elution buffer (25 mM HEPES-KOH pH7.9, 2 mM MgCl_2_, 20% glycerol, 300 mM KCl, 1 mM DTT, 1x CPI, 0.4 mg/ml 3xFLAG peptide (Sigma)) totaling 1 hr. The eluted protein was dialyzed against storage buffer (50 mM HEPES-KOH pH 7.9, 150 mM KCl, 10% glycerol, 2 mM β-mercaptoethanol) and then concentrated in a 100k MWCO centrifugal filter (Amicon). Protein concentration was determined by A_280_ (ε = 308,450 cm^-1^ M^-1^).

For expression in *S. cerevisiae*, full-length *C. neoformans* cDNA encoding Dnmt5-10xHis was cloned into the 83ν vector (Li et al., 2009), and used to transform the *S. cerevisiae* strain JEL1 (Lindsley and Wang, 1993). Starter cultures were grown overnight in SC -ura medium with 2% glucose, then used to inoculate 2 L cultures of YPGL medium (1x YEP, 1.7% lactic acid, 3% glycerol, 0.12% glucose, 0.15 mM adenine) to a starting OD_600_ of 0.03. After growth at 30°C to an OD_600_ of 1.0, expression was induced by addition of 2% galactose. After 6 hr of continued growth at 30°C, cells were harvested, washed once in 1x TBS (50 mM Tris-Cl pH 7.6, 150 mM NaCl), and snap frozen. Frozen cells were lysed in a ball mill (6x 3 min at 15 Hz), resuspended in Ni-NTA lysis buffer (50 mM NaH_2_PO_4_ pH 8, 300 mM NaCl, 10% glycerol, 10 mM imidazole, 2 mM β-mercaptoethanol, 0.02% NP40, 1x CPI), and centrifuged 20,000 x *g* for 30 min at 4°C. Lysate was bound to Ni-NTA resin in batch format for 2 hr at 4°C. The resin was washed in column format using 5 bed volumes Ni-NTA buffer followed by 10 bed volumes Ni-NTA wash buffer (same as Ni-NTA lysis buffer except 20 mM imidazole). Bound protein was eluted with 4 bed volumes Ni-NTA elution buffer (same as Ni-NTA lysis buffer except 300 mM imidazole and no NP40). Eluted protein was dialyzed against storage buffer and applied to a HiTrap Q HP anion exchange column (GE Life sciences) pre-equilibrated in buffer A (50 mM HEPES-KOH pH 7.9, 150 mM KCl, 10% glycerol, 2mM β-mercaptoethanol). Fractions were collected across a 150-1000 mM KCl gradient, and those containing Dnmt5 were pooled, concentrated, dialyzed against storage buffer, and frozen. Protein concentration was determined by A_280_ (ε = 308,450 cm^-1^ M^-1^).

### DNA methyltransferase assay

DNA oligonucleotide substrates were synthesized by IDT (Coralville, IA) (Table S2). In most cases, DNA methylation was performed in multiple turnover conditions by incubating 30 nM recombinant Dnmt5 in DNMT reaction buffer (50 mM Tris pH 8, 25 mM NaCl, 10% glycerol, and 2 mM DTT) with 5 μM DNA substrate. When indicated, ATP and MgCl_2_ were added at 1 mM, and histone tail peptides were added at 5 μM. Reactions were initiated at 23°C by addition of 4 μM H-SAM (Perkin Elmer). Aliquots were removed at indicated time points and quenched in a solution of 10 mM SAM in 10 mM H_2_SO_4_. The quenched solution was pipetted onto DE81 filter paper and air dried for 15 min. Filter papers were then washed three times in 200 mM ammonium bicarbonate (5 min each), once with water (5 min), then rinsed twice in ethanol and dried for 20 min. Filters were added to scintillation fluid (Bio-Safe NA, Research Products International Corp.), and ^3^H was detected in an LS 6500 scintillation counter (Perkin Elmer). Background signal was determined using reactions lacking Dnmt5 enzyme. A reaction was considered to yield measurable signal when its signal was >2-fold above background cpm at every time point. Background signal was typically 50-100 cpm, whereas signal for productive reactions ranged from ∼1,000 to ∼100,000. For productive reactions, rates were calculated over the first 15-20 min where reaction progress was linear and <10% of available hemimethylated sites had been acted upon. These rate values were divided by Dnmt5 concentration to obtain k_obs_. Separate experiments with varying DNA concentration were performed to confirm that each substrate was present at saturating concentration. Serial dilutions verified that ^3^H detection was linear to the background signal level.

### Electrophoretic mobility shift assay

DNA oligonucleotides labeled with 5′ Cy5 were synthesized by IDT (Coralville, IA) (Table S2) and annealed in annealing buffer (20 mM Tris pH 8, 1 mM EDTA, 50 mM NaCl), after which they were purified by polyacrylamide gel electrophoresis. Recombinant Dnmt5 was incubated with labeled DNA probe (1-3 nM in a solution of 16 mM HEPES-KOH pH7.9, 8% glycerol, 40 mM KCl, 0.02% NP40, 1.6 mM DTT) for 20 min at room temperature, and then resolved in a polyacrylamide gel (4.5% acrylamide:bis 29:1 (Bio-Rad), 1% glycerol, 1xTBE) at 4°C. To assess the effects of nucleotide analogs on DNA binding, the analogs were mixed with equal concentration MgCl_2_ and then added to DNA binding reactions at a final concentration of 1 mM. Gels were imaged using a Typhoon 9400 Imager (Amersham) and densitometry was performed using ImageJ (Schneider et al., 2012). Dissociation constants were determined from plots of fraction probe bound versus Dnmt5 concentration by fitting the equation Y = b + (m − b) * 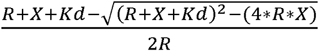 using Prism 6 (GraphPad Software) (Pagano et al., 2011). In this equation, *b* represents base signal (0, for 0% probe bound), *m* represents maximum signal (100, for 100% probe bound), *R* represents labeled probe concentration, *X* represents protein concentration, and *Y* represents percent probe bound, which was determined by quantifying the level of unbound probe.

### Generation of nucleosome substrates

DNA for nucleosome substrates was generated by PCR and corresponded to the Widom 601 nucleosome positioning sequence (147 bp) flanked on either side by a 40 bp linker sequence. Linkers each contained two CG sites that were either hemimethylated or unmethylated as dictated by the use of PCR primers containing 5mC (Table S2); the Widom 601 sequence was entirely unmethylated. Nucleosomes were assembled using purified reconstituted octamer generated with recombinant bacterially expressed histones from *Xenopus laevis.* For H3K9me3 nucleosomes, H3 histones were modified using methyl lysine analog (MLA) technology before reconstitution into octamer (Simon, 2010). Optimal ratios of DNA:octamer:dimer for nucleosome assembly were determined empirically by varying octamer:dimer in small-scale assembly reactions. Nucleosomes were reconstituted by salt dialysis over 36-48 hours, purified over 10-30% glycerol gradients, and concentrated before use.

### NADH-coupled ATPase assay

ATPase activity was assessed by incubating recombinant Dnmt5 (30-60 nM) in ATPase reaction buffer (50 mM HEPES-KOH pH 7.9, 75 mM KCl, 5% glycerol, 2 mM DTT, 2 mM phosphoenolpyruvate, 0.18 mM NADH, 0.5 mM ATP, 5 mM MgCl_2_, 10 U/ml pyruvate kinase (Sigma), and 10 U/ml lactate dehydrogenase (EMD Millipore)). DNA substrates were added typically at 5 μM in 80 μl reactions in 384 well non-stick clear bottom plates (Corning 3655). Separate experiments with varying DNA concentration were performed to confirm that each substrate was present at saturating concentration. Reactions were monitored in a Spectramax M5e plate reader (Molecular Devices) for absorbance at 340 nm and 420 nm over 30 min at 23°C. The difference between A340 and A420 was plotted versus time, and the initial linear portion of the curve was fit to determine reaction rate using Prism 6 (GraphPad Software). For measurement of *K_m,app_* ^ATP^ and *K_m,app_* ^DNA^, ATP or DNA concentration was varied in the presence of saturating amounts of DNA or ATP, respectively, and plots of rate versus concentration were fitted to the Michaelis-Menten equation.

### Protein domain identification and alignment

Identification of the Dnmt5 gene in *C. neoformans* was based on annotations of the *var. grubii* H99 genome by the Broad Institute (Cambridge, MA). Dnmt5 protein domains were identified using SMART (Schultz et al., 1998), and its primary sequence was compared to related proteins using Clustal Omega (Sievers et al., 2011).

## TABLE LEGENDS

**Table 1. ATPase kinetic parameters for Dnmt5 and Dnmt5(K1469A)**

ATPase activity was measured using 40 nM Dnmt5 in the presence of saturating concentrations of indicated DNA substrate and varying amounts of ATP. DNA substrates are described in Table S2. Values reflect average of four independent experiments and 95% confidence interval. Measurements of ATPase rate for Dnmt5(K1469A) in the presence of hemimethylated DNA and 0-1.8 mM ATP yielded poor fit to the Michaelis-Menten equation.

## SUPPLEMENTAL FIGURE LEGENDS

**Figure S1. Expression, purification, and DNA methyltransferase activity of Dnmt5**

**(A)** Coomassie stain of Dnmt5 expressed and purified in two systems. On left, *C. neoformans* Dnmt5 cDNA was cloned and expressed in *S. cerevisiae*, then purified using Ni-NTA affinity and anion exchange chromatography. On right, *C. neoformans* Dnmt5 was overexpressed from its endogenous locus using a galactose-inducible promoter, then purified using FLAG affinity resin. **(B)** dsDNA substrates used in methyltransferase experiments. Each 60 bp substrate contains three CG sites that are either unmethylated, hemimethylated, or symmetrically methylated. **(C)** Example DNA methylation kinetics using 30 nM Dnmt5 and 5 μM each of the DNA substrates in B, with or without 1 mM ATP. **(D)** Average initial rates of Dnmt5 activity on DNA substrates described in B, in the presence or absence of ATP. ND: no detected activity. Error bars represent SD; n = 3-5.

**Figure S2. Features of Dnmt5 ATPase stimulation by unmethylated and hemimethylated DNA substrates**

**(A)** Inhibition of Dnmt5 DNMT activity on hemimethylated DNA by AMP-PNP. ATP was present in all reactions at 100 μM. **(B)** Average rates of ATPase activity in the presence of 40 nM Dnmt5 and 5 μM of unmethylated DNA substrates of the indicated length. Data are normalized to ATPase rate in absence of DNA (1 min^-1^). Error bars represent SD; n = 6. **(C)** Average rates of ATPase activity in the presence of 40 nM Dnmt5 and 5 μM of unmethylated 60 bp DNA substrates with either one CG site or no CG site. Data are normalized to ATPase rate in absence of DNA (1 min^-1^). Error bars represent SD; n = 3. **(D)** Initial ATPase rates of 30 nM Dnmt5 were determined at varying concentrations of DNA substrate (unmethylated or hemimethylated) in the presence of fixed, saturating concentrations of ATP. Values were fitted to the Michaelis-Menten equation. Error bars represent SD; n = 3. **(E)** Average rates of ATPase activity in the presence of 40 nM Dnmt5 and 5 μM of DNA substrates with varying numbers of hemimethylated CG sites. Data are normalized to ATPase rate in absence of DNA (1 min^-1^). Error bars represent SD; n = 4-8. **(F)** Alignment of SNF2 domains in the same subfamily as that of Dnmt5. The Walker A and Walker B motifs are identified.

**Figure S3. Assessment of DNA binding by Dnmt5 and Dnmt5(345-747)**

**(A)** EMSA assessing binding of Dnmt5(345-747) or Dnmt5(1-150, W87A, Y90A) (0-50 nM) to 1 nM labeled hemimethylated 60 bp dsDNA. **(B)** EMSA assessing binding of Dnmt5(345-747) (200 nM) in the presence of 3 nM labeled hemimethylated DNA and excess unlabeled competitor unmethylated DNA (1 μM). **(C)** Example DNA methylation kinetics using 30 nM Dnmt5 or Dnmt5(345-747) and 5 μM of the DNA substrates described in Figure 1B. **(D)** EMSA assessing binding of Dnmt5 (67 nM) in the presence of 1 nM labeled unmethylated DNA and excess unlabeled competitor unmethylated or hemimethylated DNA (1 μM). **(E)** Screen of nucleotide analogs for effects on Dnmt5 DNA binding. Dnmt5 (67 nM) was incubated with labeled unmethylated DNA in the presence of nucleotide or nucleotide analog (1 mM), and fraction probe bound was measured by EMSA. Graph represents average and SD; n = 4.

