## Supplementary material for "ATP hydrolysis by the SNF2 domain of the ultraspecific maintenance methylase Dnmt5 drives recognition and modification of hemimethylated DNA": Table 1

**Table 1. ATPase kinetic parameters for Dnmt5 and Dnmt5(K1469A)**

| | **Dnmt5** | | **Dnmt5(K1469A)** | |
|---|---|---|---|---|
| | Unmethylated DNA | Hemimethylated DNA | Unmethylated DNA | Hemimethylated DNA |
| $K_{m,app}^{ATP}$ ($\mu$M) | 0.92 ± 0.26 | 5.9 ± 1.7 | 35 ± 9.6 | ND |
| $V_{max}$ ($\mu$M min$^{-1}$) | 0.20 ± 0.01 | 0.44 ± 0.03 | 0.21 ± 0.01 | ND |
| $k_{cat}$ (min$^{-1}$) | 5.0 ± 0.25 | 11.0 ± 0.75 | 5.25 ± 0.25 | ND |
