## Supplementary figures and images for "ATP hydrolysis by the SNF2 domain of the ultraspecific maintenance methylase Dnmt5 drives recognition and modification of hemimethylated DNA"

### Supplemental Figure S1

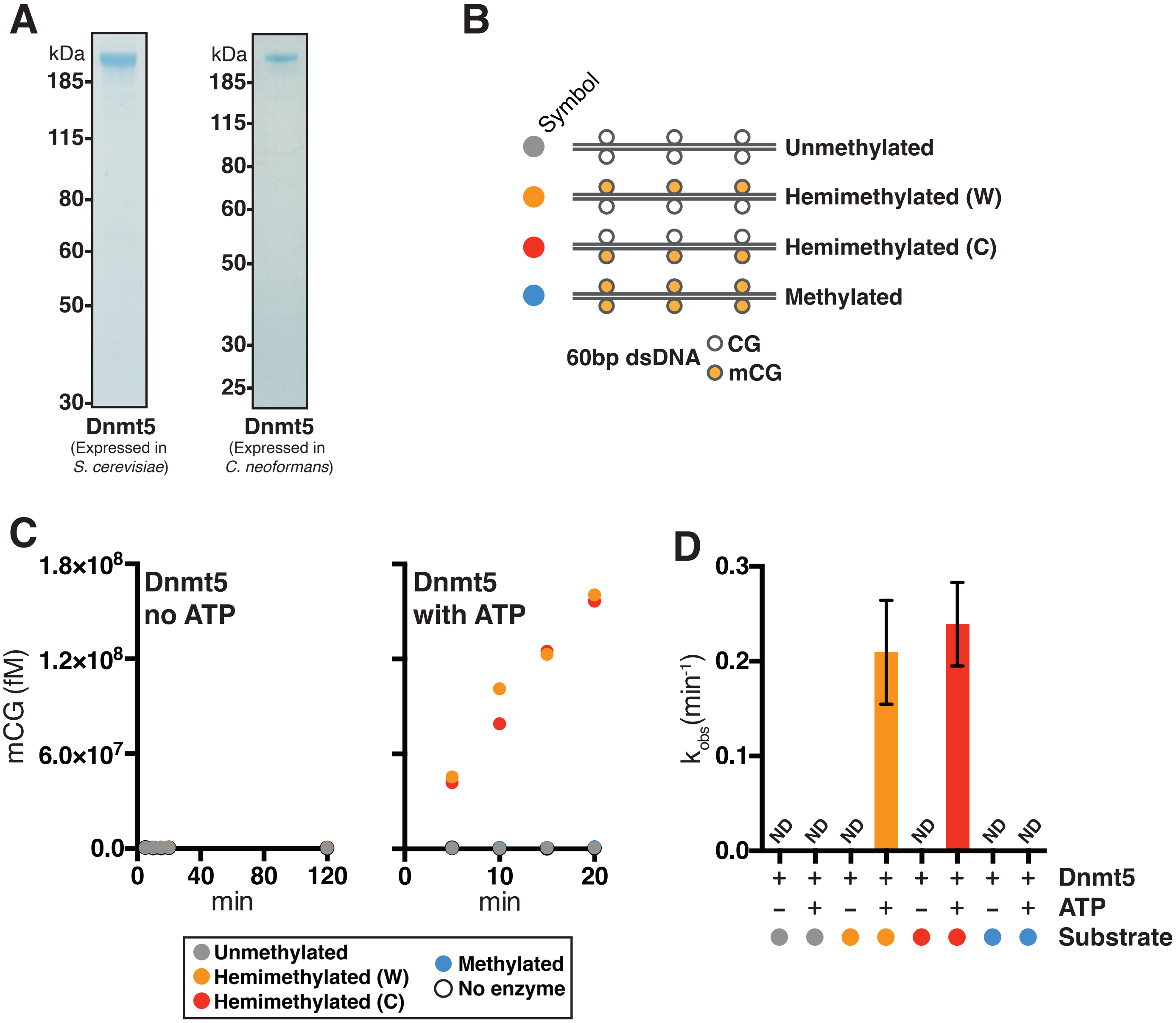

### Supplemental Figure S2

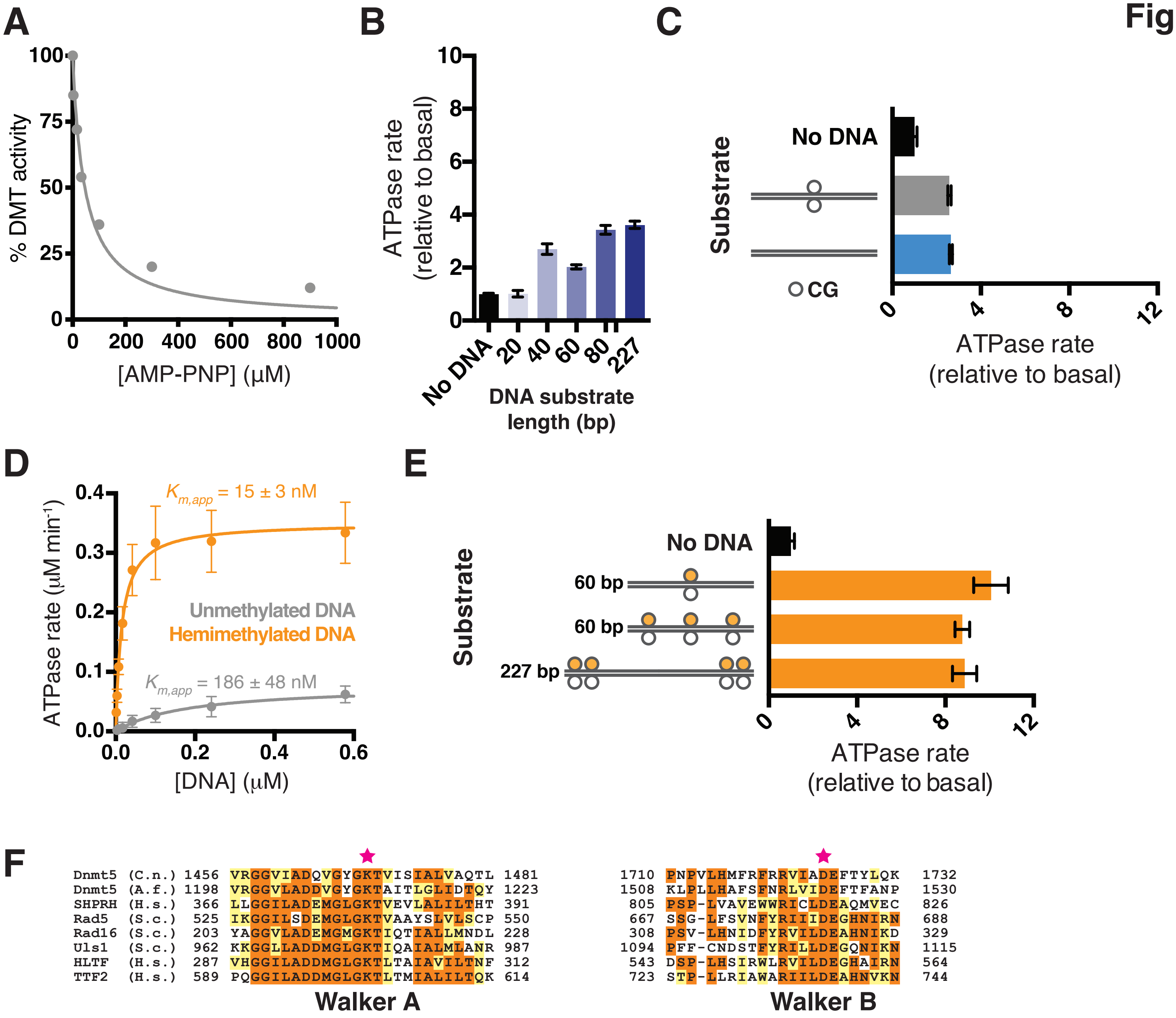

### Supplemental Figure S3

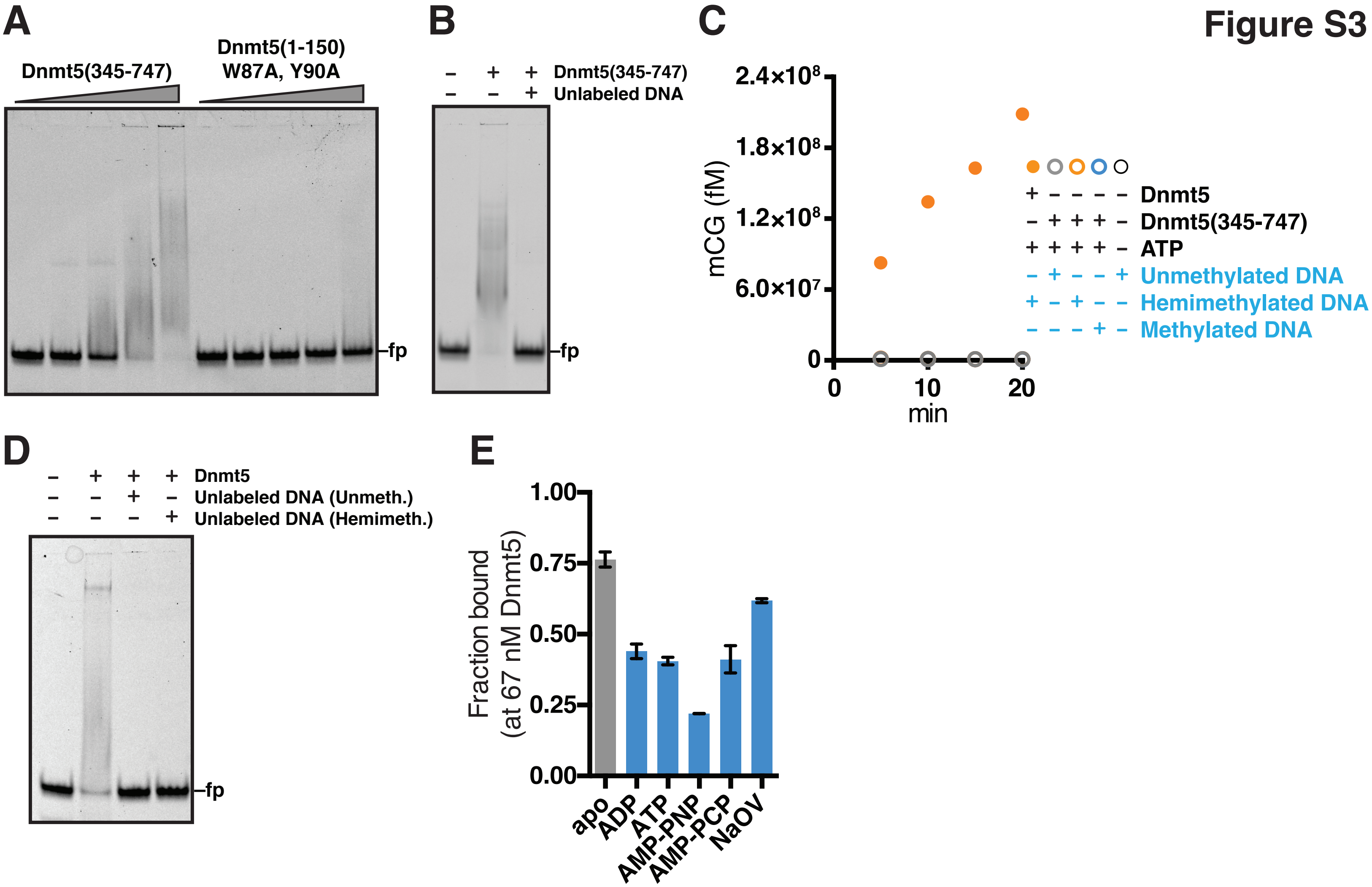
