## Supplemental Table S1 for "ATP hydrolysis by the SNF2 domain of the ultraspecific maintenance methylase Dnmt5 drives recognition and modification of hemimethylated DNA"

### Table S1. Strains Used in This Study

| PAD# | SC# | CM# | Species | Genotype | Source |
|---|---|---|---|---|---|
| 2 | - | 229 | *C. neo.* | H99 (wild-type) | 1 |
| - | 113 | - | *C. neo.* | *NeoR-pGAL7-2xFLAG-Dnmt5* | 2 |
| - | - | - | *S. cer.* | *JEL1 (α leu2 trp1 ura3-52 prb1-1122 pep4 Δhis3::PGAL1-GAL4)* | 3 |

Sources:

1= Gift of J. Lodge

2= This study

3= Lindsley and Wang. 1993 JBC 11:8096-8104
