## Supplemental Table S2 for "ATP hydrolysis by the SNF2 domain of the ultraspecific maintenance methylase Dnmt5 drives recognition and modification of hemimethylated DNA"

**Table S2. Oligonucleotides Used in This Study**

| PAD# | Name | CG sites | mCG sites | Sequence |
|---|---|---|---|---|
| 3355 | Unmethylated 60bp | 3 | 0 | CATGGCCTAAGCCGGACTGAATGAGCAAGCTTCCGGAGAATTCTGCCGGACTGCAGATGC |
| 3356 | | 3 | 0 | GCATCTGCAGTCCGGCAGAATTCTCCGGAAGCTTGCTCATTCAGTCCGGCTTAGGCCATG |
| 3354 | Hemimethylated 60bp (W) | 3 | 3 | CATGGCCTAAGC/iMe-dC/GGACTGAATGAGCAAGCTTC/iMe-dC/GGAGAATTCTGC/iMe-dC/GGACTGCAGATGC |
| 3356 | | 3 | 0 | GCATCTGCAGTCCGGCAGAATTCTCCGGAAGCTTGCTCATTCAGTCCGGCTTAGGCCATG |
| 3355 | Hemimethylated 60bp (C) | 3 | 0 | CATGGCCTAAGCCGGACTGAATGAGCAAGCTTCCGGAGAATTCTGCCGGACTGCAGATGC |
| 3371 | | 3 | 3 | GCATCTGCAGTC/iMe-dC/GGCAGAATTCTC/iMe-dC/GGAAGCTTGCTCATTCAGTC/iMe-dC/GGCTTAGGCCATG |
| 3354 | Methylated 60bp | 3 | 3 | CATGGCCTAAGC/iMe-dC/GGACTGAATGAGCAAGCTTC/iMe-dC/GGAGAATTCTGC/iMe-dC/GGACTGCAGATGC |
| 3371 | | 3 | 3 | GCATCTGCAGTC/iMe-dC/GGCAGAATTCTC/iMe-dC/GGAAGCTTGCTCATTCAGTC/iMe-dC/GGCTTAGGCCATG |
| 3352 | Solo site hemimethylated 60bp (W) | 1 | 1 | CATGGCCTAAGCaGGACTGAATGAGCAAGCTTC/iMe-dC/GGAGAATTCTGCaGGACTGCAGATGC |
| 3404 | | 0 | 0 | GCATCTGCAGTCCtGCAGAATTCTCCGGAAGCTTGCTCATTCAGTCCtGCTTAGGCCATG |
| 3342 | 1/3 sites hemimethylated 60bp (W) | 3 | 1 | CATGGCCTAAGCCGGACTGAATGAGCAAGCTTC/iMe-dC/GGAGAATTCTGCCGGACTGCAGATGC |
| 3356 | | 3 | 0 | GCATCTGCAGTCCGGCAGAATTCTCCGGAAGCTTGCTCATTCAGTCCGGCTTAGGCCATG |
| 3340 | Forward primer nuc. DNA (Unmeth.) | 2 | 0 | GAGTTCATCCCgTATGTGATGGACCCTATACGC |
| 3335 | Reverse primer nuc. DNA (Unmeth.) | 2 | 0 | TATCCGACTGGCACCGGCA |
| 3341 | Forward primer nuc. DNA (Hemimeth.) | 2 | 2 | GAGTTCATCC/iMe-dC/gTATGTGATGGACCCTATA/iMe-dC/GC |
| 3336 | Reverse primer nuc. DNA (Hemimeth.) | 2 | 2 | TATC/iMe-dC/GACTGGCAC/iMe-dC/GGCA |
| 3255 | Unmethylated 20bp | 6 | 0 | cacgcgacgcacgacgcgaa |
| 3256 | | 6 | 0 | ttcgcgtcgtgcgtcgcgtg |
| 3373 | Unmethylated 40bp | 4 | 0 | TACAATTCACTGGCCGTCGTTTTACAACGTCGTGACTGGG |
| 3374 | | 4 | 0 | CCCAGTCACGACGTTGTAAAACGACGGCCAGTGAATTGTA |
| 3375 | Unmethylated 80bp | 6 | 0 | TACAATTCACTGGCCGTCGTTTTACAACGTCGTGACTGGGAAAACCCTGGCGTTACCCAACTTAATCGCCTTGCAGCACA |
| 3376 | | 6 | 0 | TGTGCTGCAAGGCGATTAAGTTGGGTAACGCCAGGGTTTTCCCAGTCACGACGTTGTAAAACGACGGCCAGTGAATTGTA |
| 3355 | Unmethylated 60bp Cy5 | 3 | 0 | CATGGCCTAAGCCGGACTGAATGAGCAAGCTTCCGGAGAATTCTGCCGGACTGCAGATGC |
| 3370 | | 3 | 0 | /5Cy5/GCATCTGCAGTCCGGCAGAATTCTCCGGAAGCTTGCTCATTCAGTCCGGCTTAGGCCATG |
| 3354 | Hemimethylated 60bp (W) Cy5 | 3 | 3 | CATGGCCTAAGC/iMe-dC/GGACTGAATGAGCAAGCTTC/iMe-dC/GGAGAATTCTGC/iMe-dC/GGACTGCAGATGC |
| 3370 | | 3 | 0 | /5Cy5/GCATCTGCAGTCCGGCAGAATTCTCCGGAAGCTTGCTCATTCAGTCCGGCTTAGGCCATG |
| 3358 | Solo site CG/CG 60bp | 1 | 0 | CATGGCCTAAGCaGGACTGAATGAGCAAGCTTCCGGAGAATTCTGCaGGACTGCAGATGC |
| 3404 | | 1 | 0 | GCATCTGCAGTCCtGCAGAATTCTCCGGAAGCTTGCTCATTCAGTCCtGCTTAGGCCATG |
| 3352 | Solo site mCG/CG 60bp | 1 | 1 | CATGGCCTAAGCaGGACTGAATGAGCAAGCTTC/iMe-dC/GGAGAATTCTGCaGGACTGCAGATGC |
| 3404 | | 1 | 0 | GCATCTGCAGTCCtGCAGAATTCTCCGGAAGCTTGCTCATTCAGTCCtGCTTAGGCCATG |
| 3419 | Solo site mCT/CG 60bp | 0 | 0 | CATGGCCTAAGCaGGACTGAATGAGCAAGCTTC/iMe-dC/tGAGAATTCTGCaGGACTGCAGATGC |
| 3404 | | 1 | 0 | GCATCTGCAGTCCtGCAGAATTCTCCGGAAGCTTGCTCATTCAGTCCtGCTTAGGCCATG |
| 3352 | Solo site mCG/TG 60bp | 1 | 1 | CATGGCCTAAGCaGGACTGAATGAGCAAGCTTC/iMe-dC/GGAGAATTCTGCaGGACTGCAGATGC |
| 3427 | | 0 | 0 | GCATCTGCAGTCCtGCAGAATTCTCtGGAAGCTTGCTCATTCAGTCCtGCTTAGGCCATG |
| 3352 | Solo site mCG/CT 60bp | 1 | 1 | CATGGCCTAAGCaGGACTGAATGAGCAAGCTTC/iMe-dC/GGAGAATTCTGCaGGACTGCAGATGC |
| 3422 | | 0 | 0 | GCATCTGCAGTCCtGCAGAATTCTCCtGAAGCTTGCTCATTCAGTCCtGCTTAGGCCATG |
| 3411 | Solo site mCA/TG 60bp | 0 | 0 | CATGGCCTAAGCaGGACTGAATGAGCAAGCTTC/iMe-dC/aGAGAATTCTGCaGGACTGCAGATGC |
| 3427 | | 0 | 0 | GCATCTGCAGTCCtGCAGAATTCTCtGGAAGCTTGCTCATTCAGTCCtGCTTAGGCCATG |
| 3412 | Solo site XG/CG 60bp | 0 | 0 | CATGGCCTAAGCaGGACTGAATGAGCAAGCTTC/idSp/GGAGAATTCTGCaGGACTGCAGATGC |
| 3404 | | 1 | 0 | GCATCTGCAGTCCtGCAGAATTCTCCGGAAGCTTGCTCATTCAGTCCtGCTTAGGCCATG |
| 3358 | Solo site CG/XG 60bp | 1 | 0 | CATGGCCTAAGCaGGACTGAATGAGCAAGCTTCCGGAGAATTCTGCaGGACTGCAGATGC |
| 3405 | | 0 | 0 | GCATCTGCAGTCCtGCAGAATTCTC/idSp/GGAAGCTTGCTCATTCAGTCCtGCTTAGGCCATG |
| 3358 | Solo site CG/CG 60bp Cy5 | 1 | 0 | CATGGCCTAAGCaGGACTGAATGAGCAAGCTTCCGGAGAATTCTGCaGGACTGCAGATGC |
| 3426 | | 0 | 0 | /5Cy5/GCATCTGCAGTCCtGCAGAATTCTCCGGAAGCTTGCTCATTCAGTCCtGCTTAGGCCATG |
| 3412 | Solo site XG/CG 60bp Cy5 | 0 | 0 | CATGGCCTAAGCaGGACTGAATGAGCAAGCTTC/idSp/GGAGAATTCTGCaGGACTGCAGATGC |
| 3426 | | 1 | 0 | /5Cy5/GCATCTGCAGTCCtGCAGAATTCTCCGGAAGCTTGCTCATTCAGTCCtGCTTAGGCCATG |

iMe-dC = 5mC
idSp = abasic site
